# Molecular and genetic characterization of sex-linked orange coat color in the domestic cat

**DOI:** 10.1101/2024.11.21.624608

**Authors:** Christopher B. Kaelin, Kelly A. McGowan, Joshaya C. Trotman, Donald C. Koroma, Victor A. David, Marilyn Menotti-Raymond, Emily C. Graff, Anne Schmidt-Küntzel, Elena Oancea, Gregory S. Barsh

## Abstract

The *Sex-linked orange* mutation in domestic cats causes variegated patches of reddish/yellow hair and is a defining signature of random X-inactivation in female tortoiseshell and calico cats. Unlike the situation for most coat color genes, there is no apparent homolog for *Sex-linked orange* in other mammals. We show that the *Sex-linked orange* is caused by a 5 kb deletion that leads to ectopic and melanocyte-specific expression of the *Rho GTPase Activating Protein 36* (*Arhgap36*) gene. Single cell RNA-seq studies from fetal cat skin reveal that red/yellow hair color is caused by reduced expression of melanogenic genes that are normally activated by the Melanocortin 1 receptor (Mc1r)—cyclic adenosine monophosphate (cAMP)—protein kinase A (PKA) pathway, but the *Mc1r* gene and its ability to stimulate cAMP accumulation is intact. Instead, we show that increased expression of *Arhgap36* in melanocytes leads to reduced levels of the PKA catalytic subunit (PKA_C_); thus, *Sex-linked orange* is genetically and biochemically downstream of *Mc1r*. Our findings solve a comparative genomic conundrum, provide *in vivo* evidence for the ability of Arhgap36 to inhibit PKA, and reveal a molecular explanation for a charismatic color pattern with a rich genetic history.

## Introduction

Patchwork coloration in calico and tortoiseshell cats has played a central role in the conceptualization and exemplification of X-chromosome-inactivation^1^. In eutherian females, random inactivation of an X chromosome in embryogenesis is stably and clonally maintained^2^, producing a mosaic in which patches of cells express genes from only one of the two X chromosomes. Calico and tortoiseshell female cats are heterozygous for the *Sex-linked orange* (*O*) mutation—orange patches represent a clone in which the ancestral (*o*, nonorange) allele has been inactivated, with brown or black patches representing a clone in which the derived (*O*, orange) allele has been inactivated (Figure 1A). These developmental genetic events contributed to Mary Lyon’s original theory for X-inactivation in 1961^1^ but the underlying molecular cause has remained obscure.

**Figure 1:**
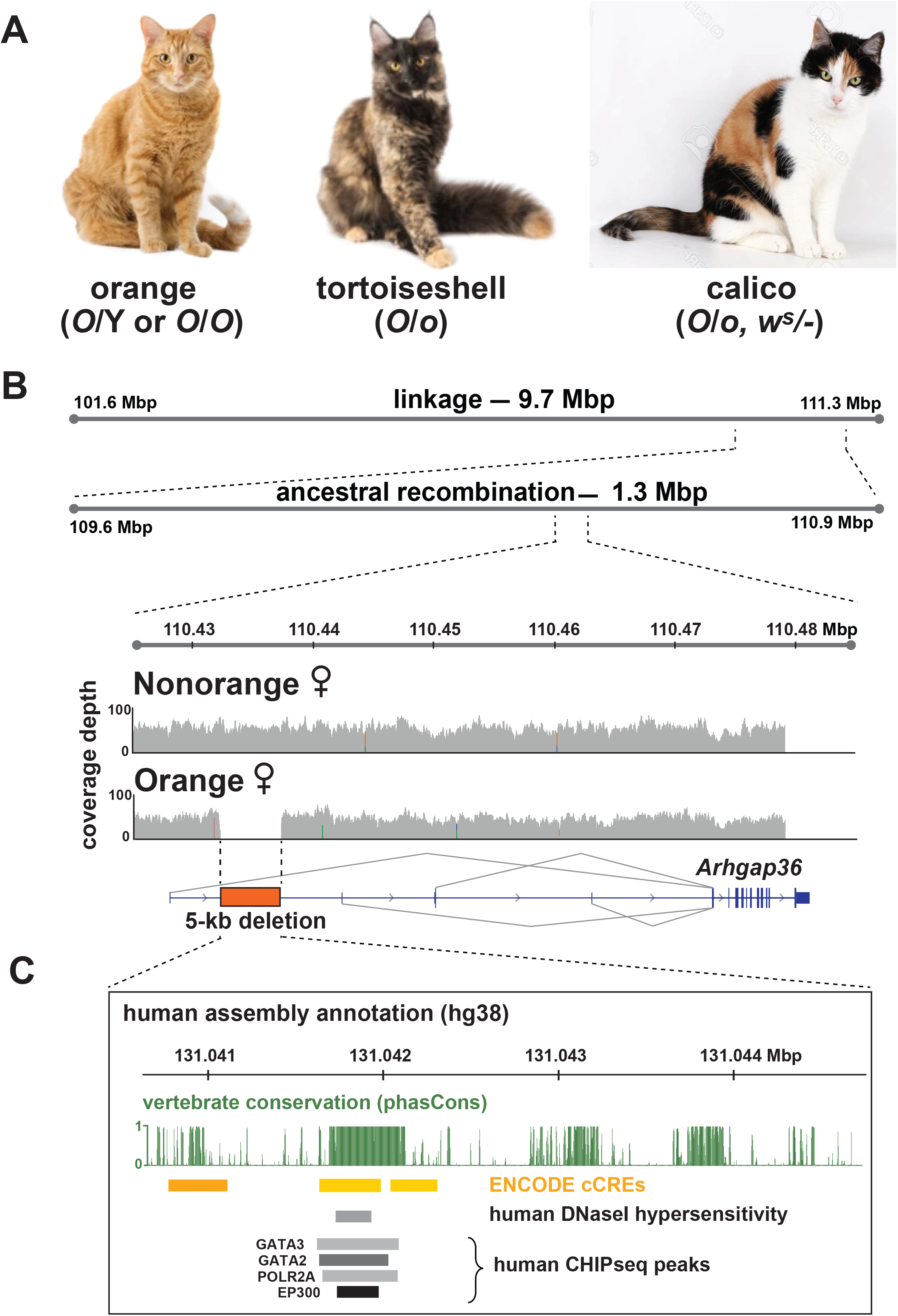
Genetic characterization of *Sex-linked orange* coat color patterns. (A) The *Sex-linked orange* mutation is responsible for orange coat color in hemizygous (*O/Y*) males and homozygous (*O/O*) females, and for patchwork color in heterozygous (*O/+*) female tortoiseshell and calico cats. In calico cats, the color patches are expanded relative to the tortoiseshell cat, caused by an autosomal dominant white spotting mutation (*w*^*s*^) at the *Kit* locus. (B) Genetic mapping of *Sex-linked orange* by linkage (A.S.K) and by ancestral recombinant breakpoints in hemizygous males (Figure S1) refines a genetic interval with a 5-kb deletion (orange box), recognized as the absence of WGS coverage (histograms) in an orange female cat and shown relative to *Arhgap36* exons (blue ticks and grey lines showing alternative promoter splicing). (C) Evolutionary conservation and gene regulatory annotation within the human interval orthologous to the *Sex-linked* orange deletion. Cat and human coordinates are from felCat9 and hg38, respectively.

The pigmentary phenotype of *Sex-linked orange* resembles that caused by loss-of-function for *Melanocortin 1 receptor* (*Mc1r*)^3^, an autosomal gene that encodes a seven transmembrane receptor expressed in melanocytes^4,5^ whose natural ligand is an antagonist, Agouti signaling protein (Asip)^6,7^. Mc1r is a Gs-coupled receptor with constitutive activity; thus, when Mc1r is expressed in melanocytes, cyclic Adenosine Monophosphate (cAMP) levels and Protein Kinase A (PKA) activity increase, while cAMP levels and PKA activity are decreased in the presence of Asip^8^. *Sex-linked orange* and *Mc1r* mutant cats^9,10^ resemble each other due to a high content of red/yellow pheomelanin and a low content of black/brown eumelanin; however *Sex-linked orange* is X-linked while *Mc1r* is autosomal. The switch between pheomelanin and eumelanin is regulated by a set of cAMP-responsive melanogenic genes including *Dopachrome tautomerase* (*Dct*), *Tyrosinase related protein 1* (*Tyrp1*), and *Premelanosome protein* (*Pmel*)^11^.

As evolutionarily conserved pigment type switching components with highly restricted functions, *Asip* and *Mc1r* are common targets for mutations that cause red/yellow color^8,12^. In contrast, sex-linked yellow or orange coat color is only observed in two species, the domestic cat^13,14^ and the Syrian hamster^15^.

Surprisingly, pedigree-based mapping studies identify distinct genetic intervals in each species, indicating different genes are involved^16,17^. The absence of similar sex-linked traits in other species raises intriguing questions about genetic mechanism and the underlying biology.

*Sex-linked orange* was one of several Mendelian coat color variants segregating in a multigenerational, domestic cat pedigree established at Nestle-Purina^18^. Schmidt-Küntzel and her colleagues mapped the locus to a 9.7 Mb genetic interval on the X chromosome^17^. In what follows we use a combination of genetic, transcriptomic, histologic, and functional assays to characterize the genetic basis and biological mechanism for orange coat color in the domestic cat. Our work confirms the essential role of PKA in pigment type-switching, demonstrates the role of a recently characterized PKA inhibitor protein, and provides a likely explanation for the absence of calico dogs, mice, and other mammals.

## Results

### Genetic mapping of *Sex-linked Orange*

Assuming *Sex-linked orange* to be a derived trait with a single, shared origin, we constructed haplotypes across the linkage interval for hemizygous orange males to delineate a 1.28 Mb interval in which all orange cats shared a common haplotype (Figures 1B, S1A, S1B). Within this interval, there were 51 variants - SNVs, indels, or structural variants - present on orange haplotypes but absent from nonorange haplotypes and the felCat9 reference genome assembly^19^. However, 48 of those 51 variants are evident in whole genome sequences (WGS) from pedigreed cats of seven breeds –Abyssinian, Bengal, Chartreux, Egyptian Mau, Ocicat, Havana Brown, and Toyger—that do not permit orange or calico coat color in the breed standard (https://cfa.org/breeds/). The three remaining variants include an SNV within a fixed SINE element (chrX:110,431,605), a nucleotide repeat expansion allele (chrX:109,896,099), and a 5,074 bp deletion (chrX:110,432,079-110,437,152).

The 1.28 Mb genetically defined interval is contiguous with no gaps or ambiguities^19^. Within this interval, the SINE-based SNV and the nucleotide repeat expansion are in non-conserved regions that lack signatures of gene regulatory function. On the other hand, the deletion lies between alternative transcriptional start sites for *Rho GTPase Activating Protein 36* (*Arhgap36*) (Figure 1B), spans an ultra-conserved mammalian noncoding element^20^, and harbors transcription factor binding sites associated with elevated *Arhgap36* expression in neuroblastomas^21,22^ (Figure 1C). The deletion is perfectly concordant with orange coat color in 188 additional cats (145 orange, 6 calico/tortoiseshell, and 37 nonorange cats, Table 1). Taken together, these observations provide strong genetic and genomic evidence that the 5 kb deletion causes *Sex-linked orange*.

**Table 1:** Association of the *X110*.*432_107*.*437del* with orange/calico coat color.

| **Color** | ***+/Y or +/+*** | ***+/del*** | ***del/Y or del/del*** |
| --- | --- | --- | --- |
| Orange | 0 | 0 | 145 |
| Calico/Tortoiseshell | 0 | 6 | 0 |
| Non-orange | 37 | 0 | 0 |

### Altered gene expression in orange cats

There are 17 annotated genes within the 1.28 Mb genetic interval and, in skin biopsies from orange and nonorange cats, expression by qRT-PCR was detectable for seven of those genes. Six genes showed no difference in expression levels; however, *Arhgap36* expression was elevated 13-fold in orange relative to nonorange skin (Figure 2A, n=4 orange and 4 nonorange cats, Student’s *t*-test *P*-value=0.013). We also measured the expression of pigmentation genes that are transcriptional targets of Mc1r signaling. Levels of *Dct, Mitf, Pmel*, and *Tyrp1* RNA are significantly reduced in orange cat skin (Figure 2B), similar to mouse models of impaired Mc1r signaling^23,24^. As described below, functional studies from our group and others^25,26^ provide a mechanistic link between *Arhgap36* overexpression and pheomelanogenesis and, in what follows, we refer to orange and nonorange individuals as *Arhgap36*^*del*^ (*Arhgap36*^*del*^/Y or *Arhgap36*^*del/del*^) and *Arhgap36*^*+*^ (*Arhgap36*^*+*^*/*Y or *Arhgap36*^*+/+*^), respectively.

**Figure 2:**
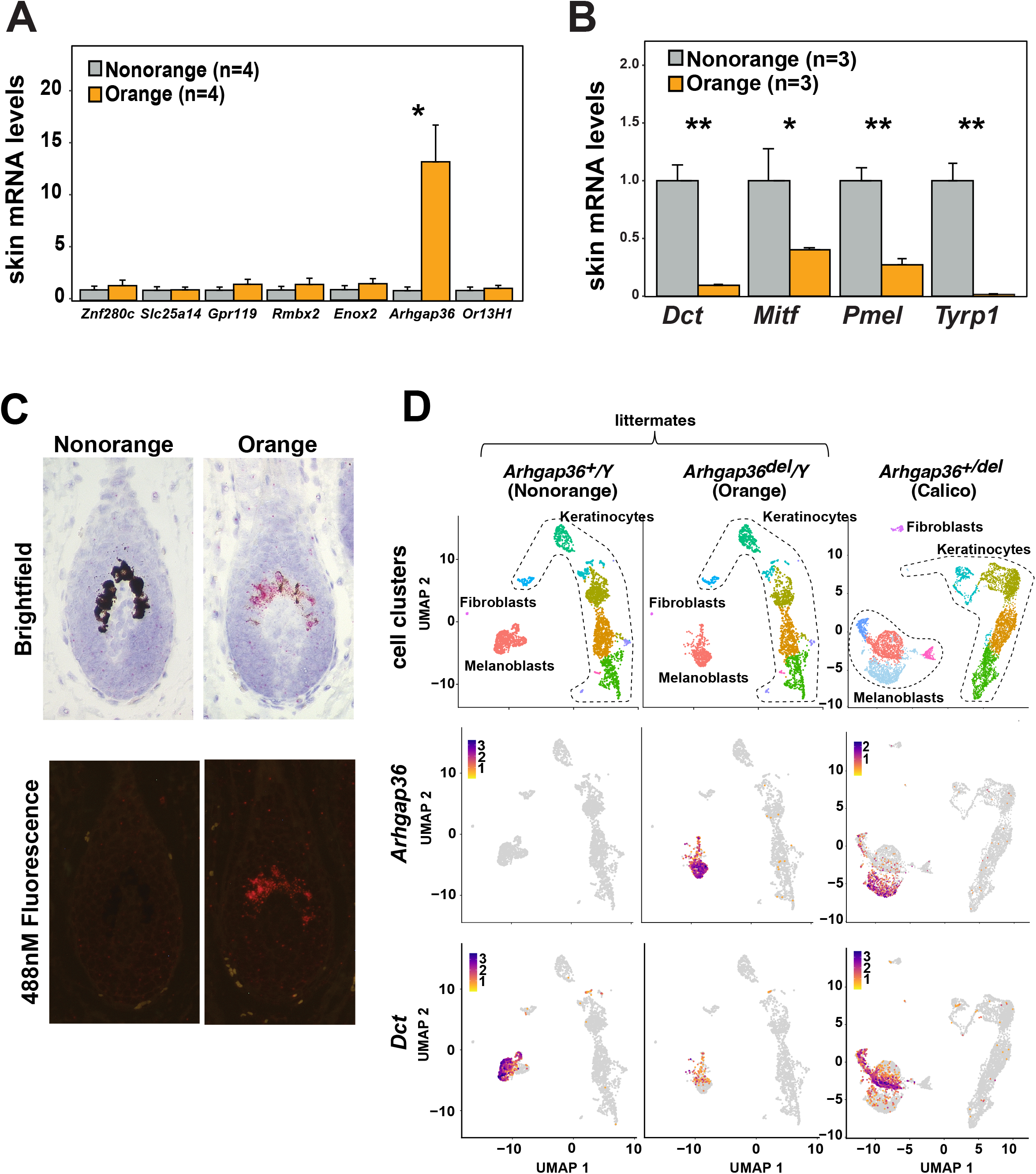
Altered gene expression in orange cats. (A) RNA levels of genes in the 1.28-Mb *Sex-linked orange* candidate interval, measured by qRT-PCR from skin biopsies of orange and nonorange adult cats (n=4 per group, \**P*=0.013, Bonferroni-adjusted Student’s t-test). (B) RNA levels of pigment cell-specific genes, measured by qRT-PCR from skin biopsies of orange and nonorange male, neonatal cats (n=3 per group, * *P*<0.10 and ** *P*<0.01, Student’s t-test). (C) Localization of *Arhgap36* RNA (pink and red, in brightfield and fluorescence images, respectively) and eumelanin (black) in histologic sections of hair bulbs from fetal skin biopsies. Fluorescence images permit visualization of *Arhgap36* RNA without potential interference from eumelanin. (D) *Top panels* – UMAP projections of single-cell RNAseq from fetal epidermal skin, colored by shared nearest neighbor clustering and annotated according to cell lineage. *Middle and bottom panels* – UMAP feature plots of *Arhgap36* and *Dct* expression, respectively.

*Arhgap36* is normally expressed in neuroendocrine organs, including the adrenal gland, hypothalamus, and pituitary gland^27^. We examined *Arhgap36* expression in these and several other tissues and observed no difference between *Arhgap36*^*del*^ and *Arhgap36*^*+*^ cats (Table S1). However, using *in situ* hybridization to examine mRNA in neonatal skin, *Arhgap36* expression was not detectable in dermis, epidermis, or hair follicles from *Arhgap36*^*+*^cats, but exhibited robust and specific expression in hair follicle melanocytes from *Arhgap36*^*del*^ cats (Figure 2C).

To further explore the relationship between gene expression and the *Arhgap36* deletion, we carried out single-cell RNA sequencing (scRNA-seq) studies of fetal skin preparations. Pigment cells are a small fraction of all skin cells but, at 44 to 48 days of gestation, developing hair follicles have yet to erupt through the skin^28^, which allows melanoblasts to be enriched from epidermal sheets using antibodies against Kit. In scRNA-seq libraries prepared from *Arhgap36*^*del*^/Y and *Arhgap36*^*+*^/Y littermates, co-clustering of both samples identified spatially coincident fibroblasts and keratinocytes populations, but spatially distinct melanoblast populations, in UMAP projections (Figure 2D). As described below, hundreds of genes are differentially expressed between *Arhgap36*^*del*^/Y and *Arhgap36*^*+*^/Y melanoblast populations, including *Arhgap36* itself (expressed at high levels in *Arhgap36*^*del*^/Y cells, middle panel, Figure 2D) and targets of Mc1r signaling such as *Dct* (lower panel, Figure 2D). Notably, *Arhgap36* expression was detectable only in melanoblasts and only from the *Arhgap36*^*del*^/Y sample.

A total of 457 genes show differential expression between *Arhgap36*^*del*^/Y and *Arhgap36*^*+*^/Y melanoblasts (adjusted *P*-value < 0.05, Wilcoxon rank sum test, and > 2-fold expression difference, Figures 2D, S2A), nearly an order of magnitude more than comparisons of equivalently sized keratinocyte populations. To refine and validate the set of differentially expressed genes between *Arhgap36*^*del*^ and *Arhgap36*^*+*^ cells, we examined a melanoblast-enriched scRNA-seq library from an *Arhgap36*^*+/del*^ female, in which X chromosome inactivation causes clonally inherited mosaicism in an otherwise isogenic background.

Clustering revealed two large (and two smaller) melanoblast populations defined by 473 differentially expressed genes (adjusted *P*-value < 0.05, Wilcoxon rank sum test, and > 2-fold expression change, Figures 2D, S2B). Because *Arhgap36* is expressed predominantly in one of the two populations (*P*-value =1.93×10^−142^, Figure 2D), we infer that the clustering is driven by whether the *Arhgap36*^*+*^ or *Arhgap36*^*del*^ bearing X chromosome was inactivated. Considering the intersection between the two experiments— comparison of male littermates, and comparison of melanoblast populations from a mosaic female—286 genes are differentially expressed, with 166 and 120 genes showing higher and lower expression, respectively, in *Arhgap36*^*del*^ melanoblasts (Figure S2C, Table S2).

Two distinct biologic functions are apparent from our scRNA-seq results. First, *Arghap36*^*del*^ cells exhibit a significantly reduced expression of pigment cell-specific genes required for eumelanin (but not pheomelanin) synthesis, including *Pmel, Oca2, Slc24a5, Tyrp1*, and *Dct* (Figure S3A). These observations provide a straightforward explanation for the similar red/yellow hair color phenotypes of orange cats and loss-of-function *Mc1r* mutants in cats^9,10^ and other mammals^12^. Second, *Arhgap36*^*del*^ cells exhibit a significant reduction in expression of *Pkib* as well as 6 genes that encode cAMP-degrading phosphodiesterases, including *Pde3a, Pde3b, Pde4b, Pde4d, Pde9a*, and *Pde10a* (Figure S3B, Tables S2 and S3). *Pkib* encodes an inhibitor of the PKA catalytic subunit (PKA_C_) and its downregulation would be expected to increase PKA_C_ activity. Likewise, downregulation of phosphodiesterases would be expected to increase intracellular cAMP levels, also increasing PKA_C_ activity. In summary, decreased expression of eumelanogenic genes that are normally activated by cAMP together with decreased expression of genes that normally reduce PKA_C_ activity suggests that the fundamental defect in *Arhgap*^*del*^ cells affects signal transduction downstream of cAMP and PKA_C_.

In the littermate and the calico datasets described above (Figure 2D), *Arhgap36* expression was both melanoblast- and *Arhgap36*^*del*^-specific. We also generated a scRNA-seq library from whole skin of an *Arhgap36*^*del*^/Y embryo at mid-gestation. We identified multiple different cell types besides melanoblasts including myoblasts, vascular and lymphatic epithelium, neural crest cells, dendritic cells, and dermal fibroblasts, but melanoblasts were the only cell type in which *Arhgap36* expression was detected (Figure S4). These results together with the absence of any transcript counts from *Arhgap36*^*+*^ melanoblasts shows that ectopic expression of *Arhgap36* in pigment cells (but not neural crest cell types) is a molecular signature for the *Sex-linked orange* mutation.

### Arhgap36 function in pigment type switching

Shortly after we discovered the *Arhgap36* deletion and associated increase in gene expression in orange cats, Eccles et al. reported that Arhgap36 functioned not as a Rho GTPase inhibitor but as an inhibitor of PKA signaling^25^ by directly binding the PKA_C_ and targeting it for lysosomal degradation. Transcriptional activation of Mc1r-regulated genes depends on nuclear phosphorylation of the Cyclic AMP-response element binding protein (CREB) transcription factor by PKA_C_^29–31^; thus, an Arhgap36-mediated depletion of PKA_C_ protein would provide a functional model connecting melanocyte-specific ectopic expression of *Arhgap36* to its effects on coat color.

Human *ARHGAP36* encodes five alternative isoforms that differ in their transcriptional initiation sites and amino-terminal sequence that helps determine intracellular localization. Three of the human isoforms (2, 4, 5) localize predominantly to the plasma membrane, which is required for inhibition of PKA_C_ ^21,25,32^. In bulk RNA sequencing of fetal epidermis from orange cats, we identified two alternative transcripts initiating from the same promoter that encode orthologs of human isoforms 4 and 5 (Figure 3A).

**Figure 3:**
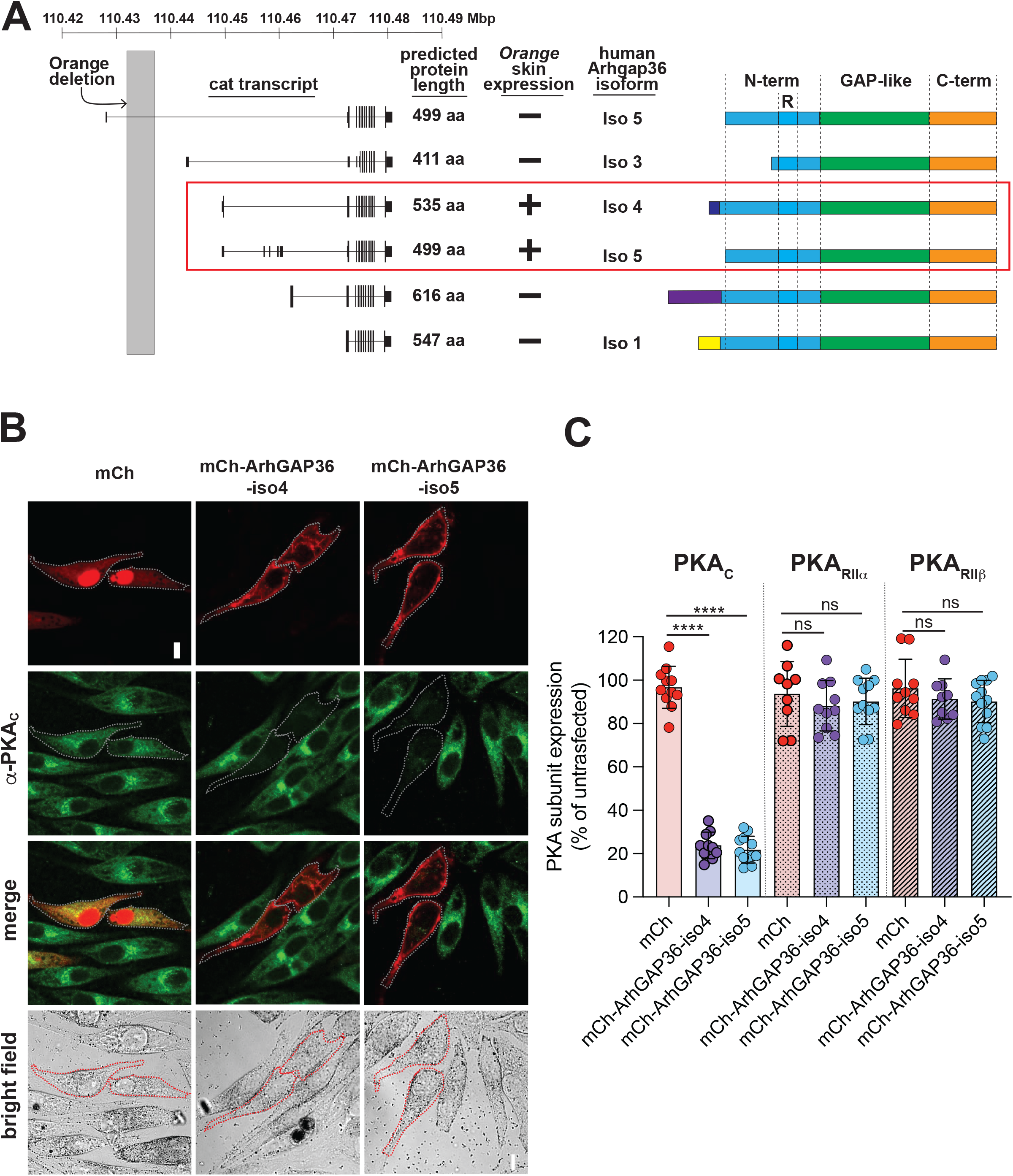
Arhgap36 expression leads to depletion of PKA_C_. (A) Diagram of cat transcripts corresponding to different human Arhgap36 isoforms and their predicted protein lengths^27, 21,25,32^. Human isoform 2 does not have a homologous cat transcript; there are two transcripts that can give rise to isoform 5, one of which begins with an untranslated first exon that lies upstream of the *Arhgap36* deletion. (B) Expression of mCh-ArhGAP36-iso4 or mCh-ArhGAP36-iso5 results in depletion of the catalytic subunit of PKA (PKA_C_). Representative confocal images of MNT1 cells expressing mCh (left column), mCh-ArhGAP36-iso4 (middle column) or mCh-ArhGAP36-iso5 (right column) and immunostained with antibodies against PKA_C_. Cells expressing mCh had similar levels of endogenous PKA_C_ immunofluorescence compared to neighboring non-transfected MNT1 cells (left column), while expression of mCh-ArhGAP36-iso4 (middle column) or mCh-ArhGAP36-iso5 (right column) led to decrease PKA_C_ immunofluorescence signal. Calibration bar: 10 μm. (C) Quantification of the average PKA_C_, PKA_RIIα_, PKA_RIIβ_ immunostaining intensity of mCh-ArhGAP36 or mCh-expressing cells calculated as the average intensity of each antibody for all cells expressing mCh compared to cells that did not express mCh in a single image. In contrast to decreased PKA_C_ expression in cells expressing both isoforms of mCh-ArhGAP36, no significant differences were observed in the expression levels of PKA_RIIα_ and PKA_RIIβ_ in MNT1 cells (see representative images in Fig. S6). Each dot represents one image, ≥50 cells/condition from n≥3 independent experiments; students T-test, **** P < 0.0001, ns not significant.

**Figure 4:**
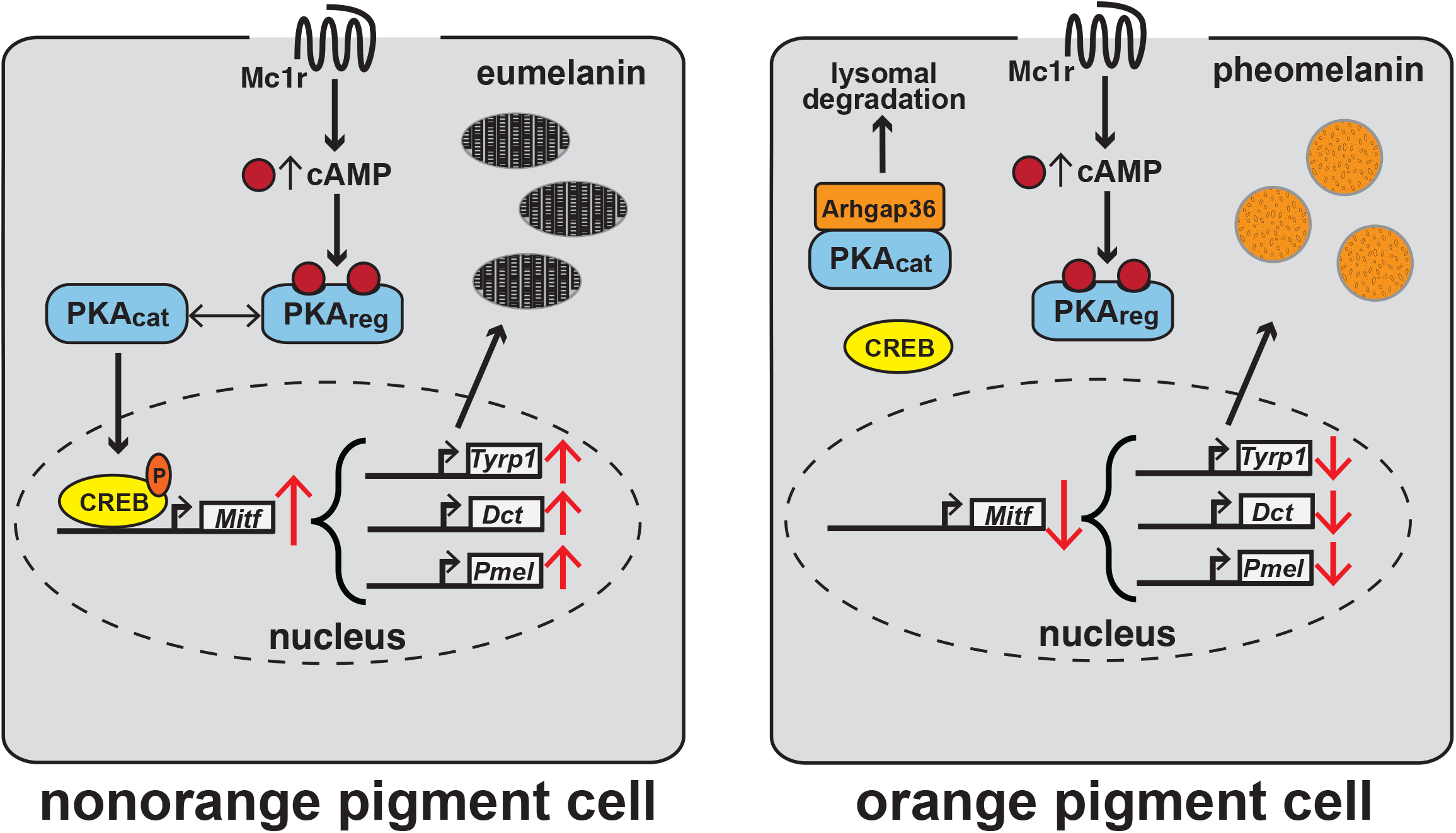
Mechanism of Arhgap36-dependent production of red/yellow pigment. In *Arhgap36*^*+*^ melanocytes (left), constitutive Mc1r activity stimulates cAMP production, allowing dissociation and nuclear translocation of PKA_C_ from the PKA apoenzyme, followed by transcriptional activation of melanogenic genes required for eumelanin production. In *Arhgap36*^*del*^ melanocytes (right), ectopic expression of Arhgap36 binds to and targets PKA_C_ for degradation, downregulating *Tyrp1, Dct*, and *Pmel*, which leads to pheomelanin production.

Previous work on Arhgap36 and PKA_C_ utilized kidney cell lines (MDCK and HEK293T cells)^25^ and motor neurons^26^. To investigate Arhgap36 function in relevant cellular context, we generated and expression constructs encoding the domestic cat *Arhgap36* isoforms 4 and 5 tagged with mCherry (mCh) at the N-terminus (mCh-ArhGAP36-iso4 and mCh-ArhGAP36-iso5) and expressed them in human melanoma MNT1 cells. Both isoforms localized at or near the plasma membrane and partially in intracellular compartments (Figure 3B). MC1R activation by α-melanocyte stimulating hormone (α-MSH) induced a larger cAMP accumulation in Arhgap36 expressing MNT1 cells compared to control (mCh) transfected cells (Figure S5). The elevated cAMP levels are consistent with the Arhgap36-associated transcriptional downregulation of cAMP-metabolizing phosphodiesterases in our scRNA-seq data and indicate that Arhgap36 attenuation of MC1R signal transduction occurs downstream of cAMP. When we used immunostaining to compare the expression of endogenous PKA in transfected MNT1 cells, and we found that levels of PKA_C_ but not PKA_R_ were significantly decreased (Figure 3B and C, S6A and B). The ability of cat Arhgap36 isoforms to deplete PKA_C_, but not perturb cAMP responsiveness in MNT1 cells is consistent with Arhgap36 function in other cellular contexts and supports a functional mechanism wherein ectopic *Arhgap36* expression attenuates MC1R signaling by inhibiting PKA, thereby dampening melanogenic genes expression and prompting a switch from eumelanin to pheomelanin synthesis.

## Discussion

The genetics of pigment type-switching has provided fundamental insight into many different cell types, tissues, and disease states, largely because key components of the melanocortin system—inverse agonists, accessory proteins, and Gs-coupled receptors that act to increase intracellular cAMP—are conserved in both structure and function across mammalian evolution^8,33^. Although the *Sex-linked orange* mutation in tortoiseshell and calico cats is an exception in that homologous mutations are not apparent in other mammals, our work reveals an underlying mechanism that is highly conserved and that lies downstream of cAMP.

Arhgap36 was named based on its sequence similarity to a family of rho GTPase activating proteins but is missing a critical zinc finger motif and lacks detectable GAP activity^34^. Instead, Arhgap36 was identified in a genome-wide screen as an activator of the hedgehog signaling pathway^21^, a role that was later shown to be mediated by inhibition of PKA signaling, through direct interaction with and subsequent lysosomal degradation of PKA_C_.^25^ *Arhgap36* knockout mice are viable and fertile but exhibit abnormalities in spinal cord patterning that is thought to involve disruption of hedgehog signaling^26^.

Increased expression of *Arhgap36* has been implicated in medulloblastoma, neuroblastoma, and several endocrine cancers^21,25,35,36^. In addition to our results on orange cats, there is one reported case of a germline gain-of-function mutation in humans that is associated with a chromosomal rearrangement, enhancer hijacking, and ectopic expression of *ARHAGAP36* in patient fibroblasts^37^. The affected individual died at 8 years of age due to severe heterotopic ossification that was thought to be caused by *ARHGAP36*-induced activation of bone formation in joints and muscle.

As shown here, increased expression of *Arhgap36* in orange cats is limited to developing and mature melanocytes, which helps to explain the absence of non-pigmentary phenotypes in orange cats that might otherwise compromise their appeal as companion animals. Besides distinctive hair color, the only documented phenotype associated with orange coat color is lentigo simplex^38^, a benign hypermelanosis and hyperplasia of epidermal melanocytes evident in glabrous regions, especially the lips, nose, eyelids, and gingiva. The lentigines resemble those typical of Carney Complex syndrome in humans, which is caused by dominant inactivating mutations in genes encoding PKA_R_ subunits^39–41^. The 5 kb deletion in orange cats includes evolutionarily conserved regulatory elements shown to bind several transcriptional activators, including JUNB/Fos/Mef2/NPAS4 in developing cortical neurons, and Tbx2, Gata2/3, and EP300 in neuroblastoma cells lines^22^. Our gene expression results from a broad series of non-pigmentary cells and tissues suggest these regulatory elements are not required for normal *Arhgap36* expression. It is possible that regulatory elements within the deletion can function as novel repressors of *Arhgap36* expression specifically in melanocytes, but a more likely explanation is that the deletion generates ectopic enhancer function by shuffling binding motifs in a genomic interval primed for enhancer activity. If so, ectopic expression of *Arhgap36* in orange melanocytes would depend on precise juxtaposition of nearby regulatory motifs and would help to explain the absence of homologous alleles in dogs, laboratory mice, and humans.

Random X-inactivation in female embryos heterozygous for the *Arhgap36* deletion underlies both the tortoiseshell and calico color patterns (Figure 1A), in which patches of red/yellow or black/brown hair represent clonal expansion of a developing melanocyte in which either the normal or deletion-bearing X chromosome has been inactivated. The difference between tortoiseshell and calico cats is the presence of an additional white spotting mutation in calico that affects the ability of developing melanocytes to survive as they migrate away from the neural crest, allowing melanocyte clones that do survive to expand in a larger body region. Indeed, that characteristic—larger patch size of a mosaic coat color mutation in the presence of white spotting—is a developmental genetic signature of a melanocyte-autonomous process^42^. This line of reasoning is consistent with our observation that *Arhgap36* ectopic expression occurs only in pigment cells; it also helps to explain why cats that are doubly mutant for *Arhgap36* and an *Mc1r* mutation known as amber exhibit an orange rather than an amber phenotype^9^.

Forward genetic studies of coat color in the laboratory mouse have significantly contributed to a detailed molecular understanding of pigment cell biology, and more generally, the pathways underlying cell migration, stem cell biology, organelle biogenesis/trafficking, and cell signaling.^43^ Similar forward genetic approaches in non-traditional genetic models, enabled by expanding genomic capacity, extend the opportunity for biologic insight.

### Methods Biological samples

DNA samples from cats used to refine *the Sex-linked orange* genetic interval and to test association with the *Arhgap36* intronic deletion include samples form Schmidt-Küntzel et al. (2009) and Kaelin et al. (2012), collected from cats in Frederick, Maryland, the Northern California Bay Area, and Port Alegre, Rio Grande do Sul, Southern Brazil^11,17^. An orange female cat from a spay-neuter clinic in Northern California (Sja149) and a nonorange female Abyssinian (Cinnamon) were used for whole-genome resequencing. The latter cat was resequenced independently and used to build the felCat9 assembly^19^.

Genomic DNA was extracted from buccal swabs or whole blood using a QIAamp mini kit (Qiagen), and from tissues (testes/ovary) by proteinase K digestion, followed by chloroform extraction and ethanol precipitation.

Fetal skin tissues for single-cell RNA-seq and *Arhgap36* in situ hybridization were harvested from incidentally pregnant cats at spay-neuter clinics in Northern California. Additional tissue biopsies for gene expression studies were collected from cats euthanized for reasons unrelated to this study, at Auburn University, College of Veterinary Medicine and animal shelters in Alabama and Northern California.

Tissue preparation for single-cell RNA-seq, *in situ* hybridization, and qPCR are described in relevant sections below.

All animal work was conducted in accordance with relevant ethical regulations and under a protocol approved by the Stanford Administrative Panel on Laboratory Animal Care (APLAC-9722).

### Genotyping

The genetic interval from Schmidt-Küntzel et al.^17^ was refined by Sanger genotyping of 65 SNVs discovered in Montigue et al.^44^ (Figure S1A) and in our resequencing data (Figure S1B). All PCR primer sets used for genotyping are provided in Table S4.

The *Arhgap36* intronic deletion polymorphism was genotyped with two PCR reactions, using a common forward primer upstream of the deletion, and reverse primers located within or downstream of the deletion. Genotypes were assigned based on the presence or absence of the appropriately sized amplicon on an agarose gel. Sex was determined by PCR of a Y chromosome specific *Sry* amplicon. Genotyping primer sets are provided in Table S5.

### Candidate interval refinement, whole genome resequencing, and variant detection

We analyzed haplotypes in *O*/*Y* male cats to identify ancestral recombination breakpoints that could narrow the genetic interval. By genotyping 27 male cats (17 orange and 10 nonorange) at 41 SNV loci, we observed that all orange cats were monomorphic at three markers that were polymorphic in the nonorange cats (Figure S1A), which delineated a 1.4 Mb genetic interval (Figure S1A).

To identify all orange-associated variants within this interval, we generated high coverage WGS from two female cats, one orange (*O/O*) and one nonorange (*o/o*). Truseq PCR-free whole genome sequencing libraries (Illumina) were constructed using 1 ug of genomic DNA. Each library was sequenced on four lanes of an Illumina HiSeq2500 sequencer to generate 125 nucleotide paired-end reads, which were mapped to the felCat9 genome assembly with BWA-MEM^45^, after trimming with CutAdapt^46^. Coverage depths for the orange and nonorange libraries was 45.1x and 51.9x, respectively. Single-nucleotide and short insertion/deletion variants were identified with GATK^47^, Freebayes^48^, and Bcftools^49^ variant callers. We used Lumpy^50^ to detect structural variation in the *Sex-linked orange* genetic interval which were then verified with the Integrative Genome Viewer (IGV, Broad Institute).

From the WGS data, we identified 24 SNVs within the 1.4 Mb interval; additional haplotype analysis of six male orange cats (Figure S1B) narrowed the candidate interval to 1.28 Mb. Within this region, 51 variants—SNVs, indels, and/or structural variants—distinguished the *O/O* from the *o/o* data; of these variants, 48 were found in breeds that do not permit orange or calico coat color, leaving three as candidates for the orange causal variant.

### Quantitative RT-PCR

Dorsal skin biopsies from eight adult cats (four orange and four non-orange), preserved in RNAlater (ThermoFisher), were homogenized with a Kinematica polytron rotor-stator, PT 10-35 GT in TRIzol (ThremoFisher). Biopsies from skin, liver, kidney, heart, adrenal gland, pituitary gland, and hypothalamus were harvested from six cats (three orange and three black), euthanized 28-35 days after birth. Tissues preserved in Qiazol were homogenized using a handheld rotor-stator. In all cases total RNA was isolated using a Purelink mini columns (ThermoFisher) after on-column DNAseI treatment.

For each sample, 500ng of total RNA was reverse transcribed with Superscript IV (ThermoFisher) using oligo dT and random hexamer primers. Quantitative PCR was performed with a LightCycler 1.2 (Roche) using LightCycler FastStart DNA Master SYBR Green reagents (Roche), using primer sets in Table S5. *Gapdh* served as a reference gene for relative quantification.

### Single-cell RNA sequencing and analysis

Prenatal staging was based on embryonic crown-rump length over the course of the domestic cat’s 63-day gestation, as described in previous work^28^. Skin dissociations were performed on four cat embryos. One embryo was at a mid-gestation stage (4cm C-R length, 28-32 days post coitus), when hair follicle development in the skin initiates. The remaining three embryos were at a late gestation stage (8-9 cM C-R length, 44-48 days post coitus), when nascent hair is formed but has yet to emerge from the skin, facilitating dermal-epidermal separation. The mid-gestation embryo was an orange male (*Arhgap36*^*del*^/Y). The late gestation embryos consisted of orange and nonorange male littermates (*Arhgap36*^*del*^/Y and *Arhgap36*^*+*^/Y) and a calico female (*Arhgap36*^*del*/*+*^).

Single cell preparations from the mid-gestation embryo were performed following a previously developed workflow, designed to enrich for basal keratinocytes but which also captures melanoblasts. Briefly, embryos were treated with Dispase (Stem Cell Technologies) and skin was removed by scraping with a No.15-blade scalpel. Isolated skin was dissociated with trypsin (Gibco) and red blood cells were removed with a red blood cell lysis solution (Miltenyi Biotec). Dissociated cells were incubated with at anti-CD49f antibody conjugated with phycoerythrin (ThermoFisher Scientific, clone NKI-GoH3 12-0495-82) and the antibody-bound fraction was captured after incubation with anti-phycoerythrin beads and enriched using a column-magnetic bead separation procedure (Miltenyi Biotec).

Single cell preparations from the late gestation embryo followed a modified protocol to enrich for melanoblasts. Dorsal skin was excised with micro scissors and treated overnight in Dispase (Stem Cell Technologies). Epidermal sheets, comprised of keratinocytes and migrating melanoblasts, were isolated by gentle scraping with a No.15-blade scalpel and dissociated with trypsin. Red blood cells were removed with a red blood cell lysis solution (Miltenyi Biotec). To enrich for melanoblasts, ten million dissociated cells were incubated with a phycoerythrin-conjugated, CD117 (c-Kit) Monoclonal Antibody (clone 2B8, eBioscience, ThermoFisher). The antibody bound fraction was then captured using a column-magnetic bead separation procedure (Miltenyi Biotech). In each case, the bound cell fraction was ∼200,000 cells, or ∼2% of the unenriched population.

3’ end, single-cell barcoded libraries were prepared from cell suspensions using 10x Genomics single-cell RNA sequencing platform for targeted capture of 6000 cells, using chemistry v2.0 and v3.0 for mid-gestation and late gestation libraries, respectively. Each library was sequenced on a single Illumina HiSeqX lane to generate ∼450 million reads (Table S6), which were aligned to the domestic cat fca126 genome assembly (GCF_018350175.1 with NCBI *Felis catus* Annotation Release 105) using the 10x Genomics CellRanger software (v7.1.0). The ‘cellranger count’ pipeline was used to output feature-barcode matrices. The ‘cellranger aggr’ pipeline was used to aggregate data from the orange and nonorange male littermate (*Arhgap36*^*del*^/Y and *Arhgap36*^*+*^/Y) libraries into a combined matrix.

Cell clustering and differential expression analysis were performed using Seurat (v5.0.0)^51^. Gene expression matrices were filtered to retain cells with (a) >200 expressed features (genes), (b) between 2,000 and 100,000 UMIs counts (unique transcripts), and (c) < 10% of total expression from mitochondrial genes. Data normalization, scaling, and variance stabilization were performed with SCTransform, and PCA and UMAP embedding were used for dimensionality reduction. A Wilcoxon rank sum test for differential expression was performed between melanoblast populations from male littermates (Fig. S2A) or between the two largest melanoblast populations from the female calico sample (Fig. S2B). Melanoblast gene expression levels were aggregated by sample ID (male littermates) or by cluster ID (female calico) to convey normalized, relative expression values across genes as transcripts per million transcripts (TPM, Table S2).

### RNA sequencing for *Arhgap36* isoform expression analysis

An unenriched aliquot of 2.5 million dissociated epidermal skin cells, isolated from the same late gestation stage orange male (*Arhgap36*^*del*^/Y) used for scRNA-seq, were resuspended in 200 ul of TRIzol (ThremoFisher). Total RNA was isolated using a Directzol RNA mini column (Zymo Research) and used to construct a strand-specific RNA sequencing library (New England Biolabs). The library was sequenced to generate 52 million 150 nucleotide paired-end reads on a partial lane of an Illumina Novaseq sequencer. Reads were aligned to the felCat9 assembly using STAR v.2.7.5^52^. Expression of *Arhgap36* isoforms (Figure 3A) were determined by counting read pairs that overlap isoform specific exons.

### In situ hybridization

Fetal skin from near-term (stage 21-22^28^) orange and black littermates (N=8 biologic samples, 1 orange and 1 black sample from four litters) was dissected and fixed within 12 hours of the spaying surgery.

Tissue genotypes were confirmed by PCR.

5 mM paraffin embedded sections were processed for RNA *in situ* detection using RNAScope^53^ 2.5 HD Detection Reagent – RED according to manufacturer’s instructions (Advanced Cell Diagnostics).

*Arhgap36* probes were designed to target 255-1205bp of XM_006944007.5. All photomicrographs are representative of at least ten anagen follicles from each biological replicate and were captured on a Leica DMRXA2 system with DFC5500 camera and LASV4 (v.4.2.0) software.

### Arhgap36 expression constructs and transfection

Two Arhgap36 isoforms expressed in orange cat skin (XM_006944008.4 and XM_019824288.3) and corresponding to human isoforms 4 and 5 (Figure 3A) were cloned in the pcDNA4/TO vector (Invitrogen) containing the mCherry coding sequence in AflII/HindIII restriction sites. Arhgap36 isoforms were inserted after the mCherry coding sequence using a GAGA linker. The constructs were generated and sequence-verified by Genscript.

MNT1 cells were grown in DMEM supplemented with 18% FBS, 10% AIM-V, 100 units/ml Penicillin-Streptomycin at 37 °C and 5% CO_2_. For the immunostaining experiments cells were plated on acid washed and poly-L-Lysine-coated #1 glass coverslips (Neuvitro, GG-12) at ∼70% density in a 6-well plate and incubated at 37 °C overnight. Cells were transfected using PolyMag Neo Magnetofection Reagent (OZ Biosciences) using 1 μg of plasmid DNA and 1 μl Polymag Neo solution for each well.

Cells were fixed for immunostaining 14-18 h post-transfection.

### Immunostaining, Imaging, and Image analysis

MNT1 cells were fixed in 4% Paraformaldehyde (Sigma Aldrich) for 20 min at room temperature and washed with PBS. Cells were incubated with primary antibodies diluted in blocking solution consisting of PBS with 0.2% (wt/vol) saponin (Acros), 0.1% (wt/vol) BSA (Fisher Bioreagents), 0.02% (wt/vol) sodium azide (Sigma Aldrich) for 30 min at RT, then washed with PBS 3 × 5 min, as previously described (Calvo et al., 1999). Primary antibodies used were: mouse monoclonal anti-PKA_C_ (BD Biosciences, 610981, 1:100), mouse monoclonal anti-PKA_R_IIa (BD Biosciences, 612243, 1:100) and mouse monoclonal PKA_R_IIb (BD Biosciences, 610626, 1:100). For visualization, cells were incubated for 30 min at RT with anti-mouse secondary antibodies (goat or donkey) conjugated with Alexa Fluor 488 diluted in blocking solution (1:1,000). Cells were washed 3 × 5 min with PBS and mounted with VECTASHIELD antifade mounting medium (Vector Laboratories) onto glass slides. The immunostained cells were imaged using an Olympus FV3000 confocal microscope.

Confocal images were analyzed using ImageJ software. Brightness and contrast were adjusted to visualize MNT1 cells expressing low levels of mCh-Arhgap36 or mCh (used as control). Every transfected cell was selected for comparison with 10 wild type cells in each image. For each image we calculated the average fluorescence intensity of the mCh-Arhgap36 or mCh expressing cells as a fraction of the average intensity of the wildtype (non-transfected cells). All the images from one transfection were averaged. Every transfection was equivalent to one experiment.

### cAMP imaging

The FRET-based genetically encoded cAMP indicator EPAC-S^H187^ (Epac H187) was a generous gift from the Jalink Laboratory (Netherlands Cancer Institute). Cells were transfected with Epac H187 and ∼24 h after transfection cells were serum-starved in OPTI-MEM (ThermoFisher Scientific) for another ∼24 h.

Coverslips were transferred to an imaging chamber with Ringer’s solution. Sequential fluorescence images were acquired with MetaMorph software on an inverted microscope every 10 s using CFP and FRET filter cubes: λ_ex_ = 430 nm and CFP and YFP emissions were detected simultaneously using 470±20 nm and 530±25 nm band-pass filters. After acquiring 18 baseline images (3 min), 1 μM NDP-αMSH (Sigma-Aldrich) was added, followed by a mix of 25 μM forskolin (FSK, Sigma-Aldrich) and 100 μM 3-isobutyl-1-methylxanthine (IBMX, Sigma-Aldrich) added to elicit maximal cAMP response, used for normalization. Fluorescence emission intensities were quantified as F=F_CFP_/F_YFP_. Normalized fluorescence intensities were quantified using F_norm_(t) = (F_cell_(t)− F_min_)/(F_FSK+IBMX_ − F_min_), where F_cell_ is the fluorescence of an intracellular region of interest, F_FSK+IBMX_ is the maximal fluorescence with FSK and IBMX, and F_min_ is the baseline fluorescence before stimulation. Light-induced changes in fluorescence intensity were quantified using MetaMorph and Excel software (Microsoft). NDP-αMSH, FSK and IBMX were solubilized in DMSO (Sigma-Aldrich) at >100x the final concentration, so that the final DMSO concentration in the imaging chamber remained < 1% (v/v) for all experiments.

## Acknowledgments

The authors thank Hermogenes Manuel for technical assistance and Leslie Lyons and 99 Lives Consortium for generating and organizing cat genomic resources. This research has been supported in part by the National Institutes of Health (R01AR067925) and by the HudsonAlpha Institute for Biotechnology.

## Author contributions

Conceptualization, C.B.K., K.A.M., E.O., and G.S.B.; Methodology, C.B.K., K.A.M., D.C.K., J.T.;

Formal analysis, C.B.K., K.A.M., E.O.; Resources: C.B.K., V.A.D., M.M-R., A.S-K., and E.C.G.; Data

Curation, C.B.K.; Writing, C.B.K., E.O., and G.S.B.; Visualization, C.B.K. and K.A.M.

