## Supplementary material for "Molecular and genetic characterization of sex-linked orange coat color in the domestic cat": Supplmental Tables

**Figure S1: Refinement of the *Sex-linked orange* genetic interval by recombinant breakpoint mapping.** (A) Haplotypes across the 9.7 Mb linkage interval (Figure 1), from 17 orange and 10 nonorange male cats. Individuals and SNV loci are arranged in rows and colulmns respectively. 41 SNVs are colored by major (yellow) and minor (blue) allele frequency in Orange cats. The IDs of cats with recombinant breakpoints that define a minimal genetic interval (red box) are colored red, as are the breakpoint-defining SNV coordinates. Asterisks indicate individual cats selected for additional haplotype analysis in (B). (B) Haplotypes across the 1.4 Mb interval defined in (A) for six orange male cats, including three cats whose recombination breakpoints refined the interval in (A). 24 SNVSs were identified from whole genome resequencing of an orange and nonorange female cat.

**Figure S2: Differential gene expression between orange and nonorange melanoblast populations.** UMAP projections (left panels) and expression heat maps for top DEGs (right panels) after reclustering integrated orange and nonorange male littermate (A) or calico (B) melanoblast scRNAseq populations from Figure 2D. Melanoblast populations are colored by orange/nonorange littermate (A) or by shared nearest neighbor clustering (B). 457 and 473 DEGs (adjusted *p*-value < 0.05 and > 2-fold expression difference) were identified between the populations in (A) and (B), respectively. (C) Venn diagram displaying the intersection of melanoblast DEGs from orange/nonorange male littermates populations (A, blue) or calico populations (B, red). A complete list DEGs represented in both data sets is provided in Table S4.

**Figure S3: Gene expression levels of melanogenesis and Mc1r signal transduction components in scRNAseq data sets.** Violin plots of normalized, single-cell expression from respective melanoblast populations, defined in Figure S2, for genes involved in (A) melanogenesis and (B) cAMP/PKA signaling transduction. Horizontal bars indicating median normalized expression for each population. Significance cutoffs for differential expression are: n.s. – not significant, * adjusted $P<0.05$, ** adjusted $P<1\text{x}10^{-5}$, ** adjusted $P<1\text{x}10^{-10}$, Wilcoxon rank sum test.

**Figure S4: Arhgap36 expression in mid-gestation embryonic skin.** (A) A Single-cell RNAseq UMAP projections of skin cell populations, colored by shared nearest neighbor clustering, from a mid-gestation orange (*Arhgap36$^{del}$*/Y) male embryo. (B) A feature plot and (C) violin plot indicating that normalized expression of *Arhgap36* is restricted to the melanoblast population.

**Figure S5: Expression of mCh-ArhGAP36-iso4 or mCh-ArhGAP36-iso5 results in increased cAMP production in response to a-MSH stimulation of MC1R.** MNT1 cells expressing the FRET-based cAMP indicator, Epac-H187 and mCh-ArhGAP36-iso4, mCh-ArhGAP36-iso5, or mCh alone (as control) were stimulated with 1 μM a-MSH. The a-MSH-evoked cAMP response of individual cells was monitored as the ratio of CFP and YFP fluorescence intensities and represented as a function of time. Forskolin (FSK) and IBMX were added to elicit a maximal cAMP response, used for normalization. The cAMP response elicited in cells expressing either isoform of the mCh-ArhGAP36 (red and orange traces) had a higher amplitude compared to mCh alone (gray trace) (n=3 independent experiments, mean± SEM). A.U., arbitrary units.

**Figure S6: Expression of mCh-ArhGAP36-iso4 or mCh-ArhGAP36-iso5 does not alter the expression of PKA regulatory subunits ($PKA_{RIIa/b}$).** (A-C) Representative confocal images of MNT1 cells expressing mCh (top rows), mCh-ArhGAP36-iso4 (middle rows) or mCh-ArhGAP36-iso5 (bottom rows) andf immunostained with antibodies against endogenous $PKA_{RIIa}$ (A) or $PKA_{RIIb}$ (B). No significant differences were observed in the expression levels of $PKA_{RIIa/b}$ (A and B) in MNT1 cells transfected with mCh (top rows), ArhGAP36-iso4 (middle rows), or ArhGAP36-iso5 (bottom rows). Calibration bar: 10 mm.

### Supplementary Figure 1

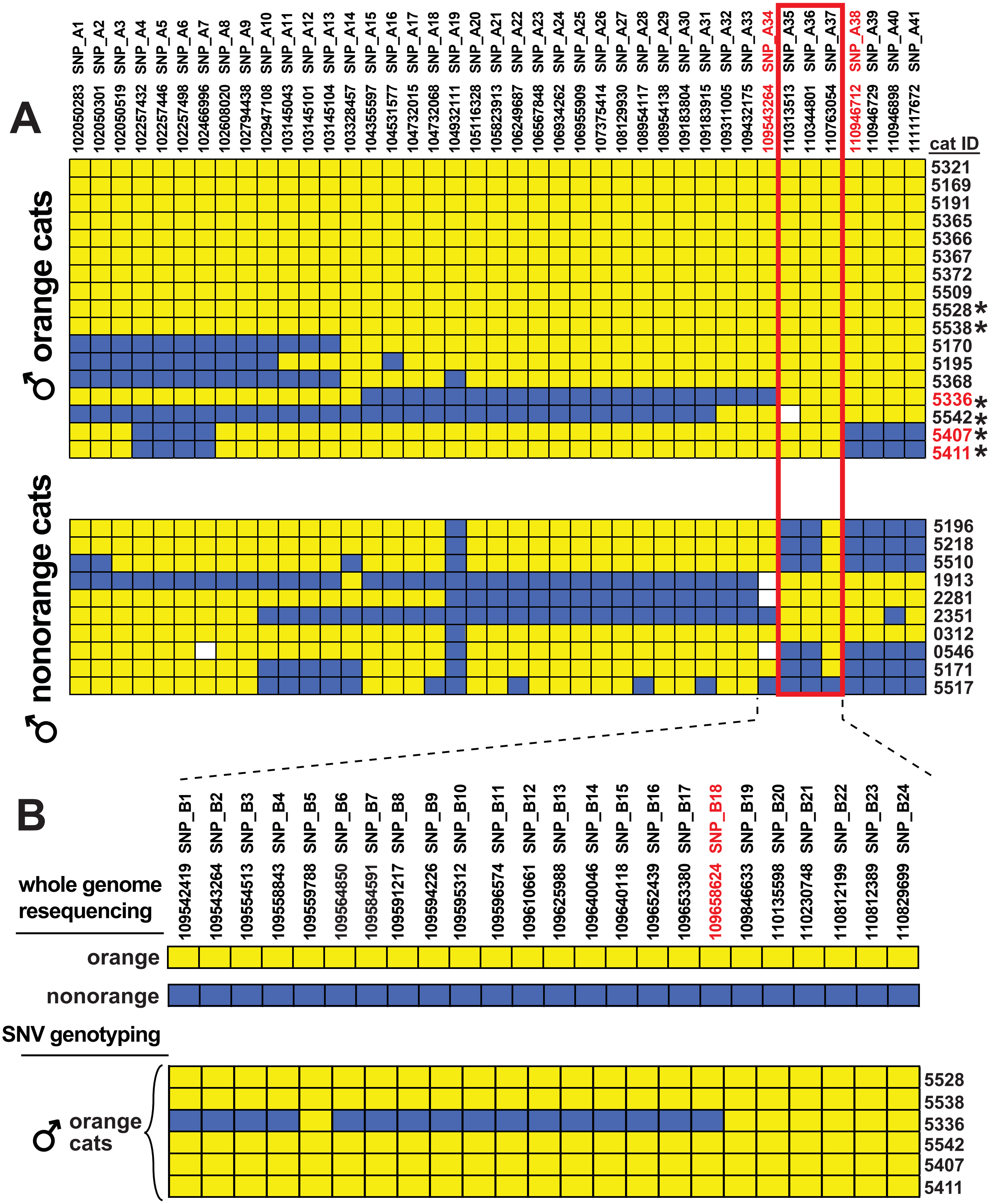

### Supplementary Figure 2

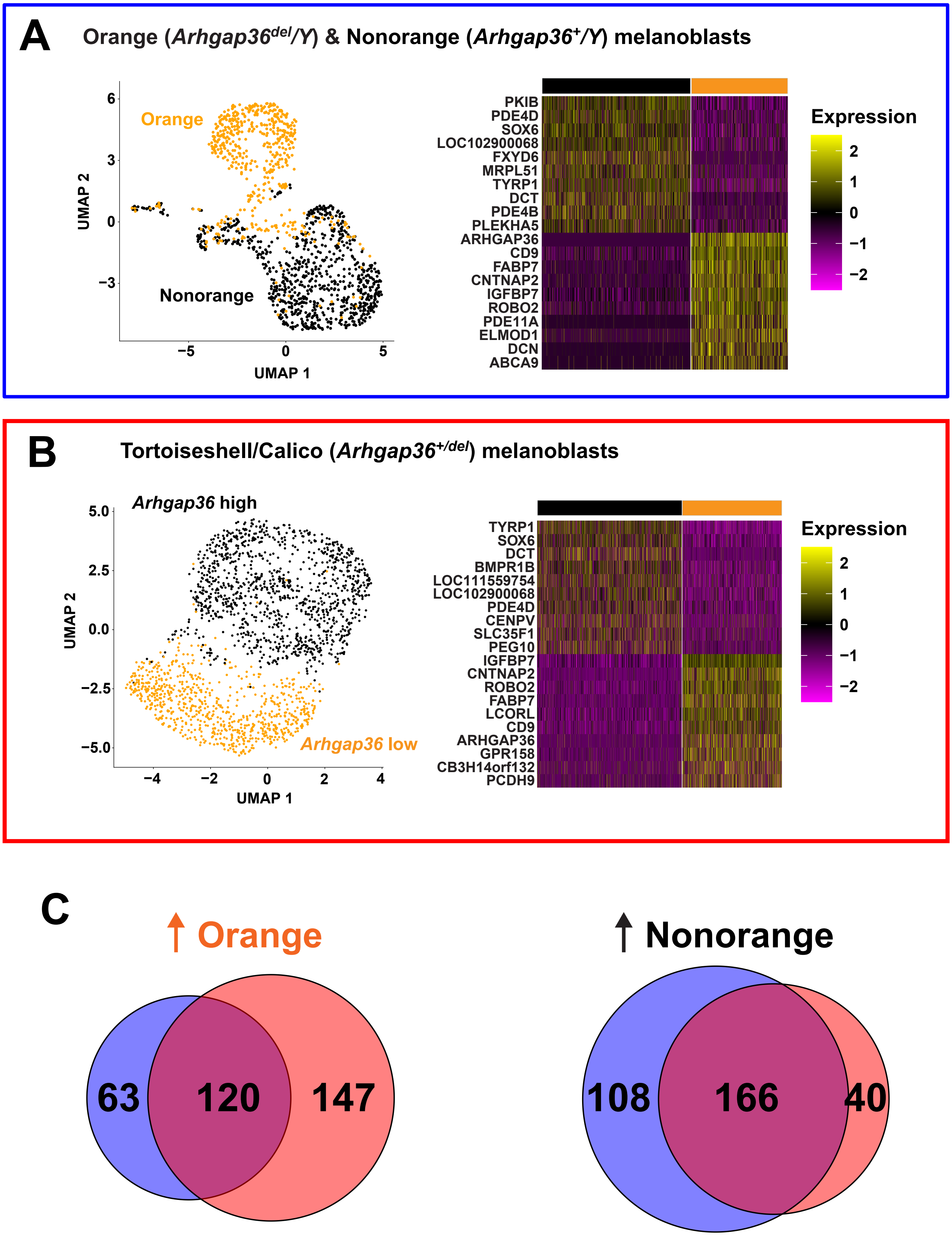

### Supplementary Figure 3

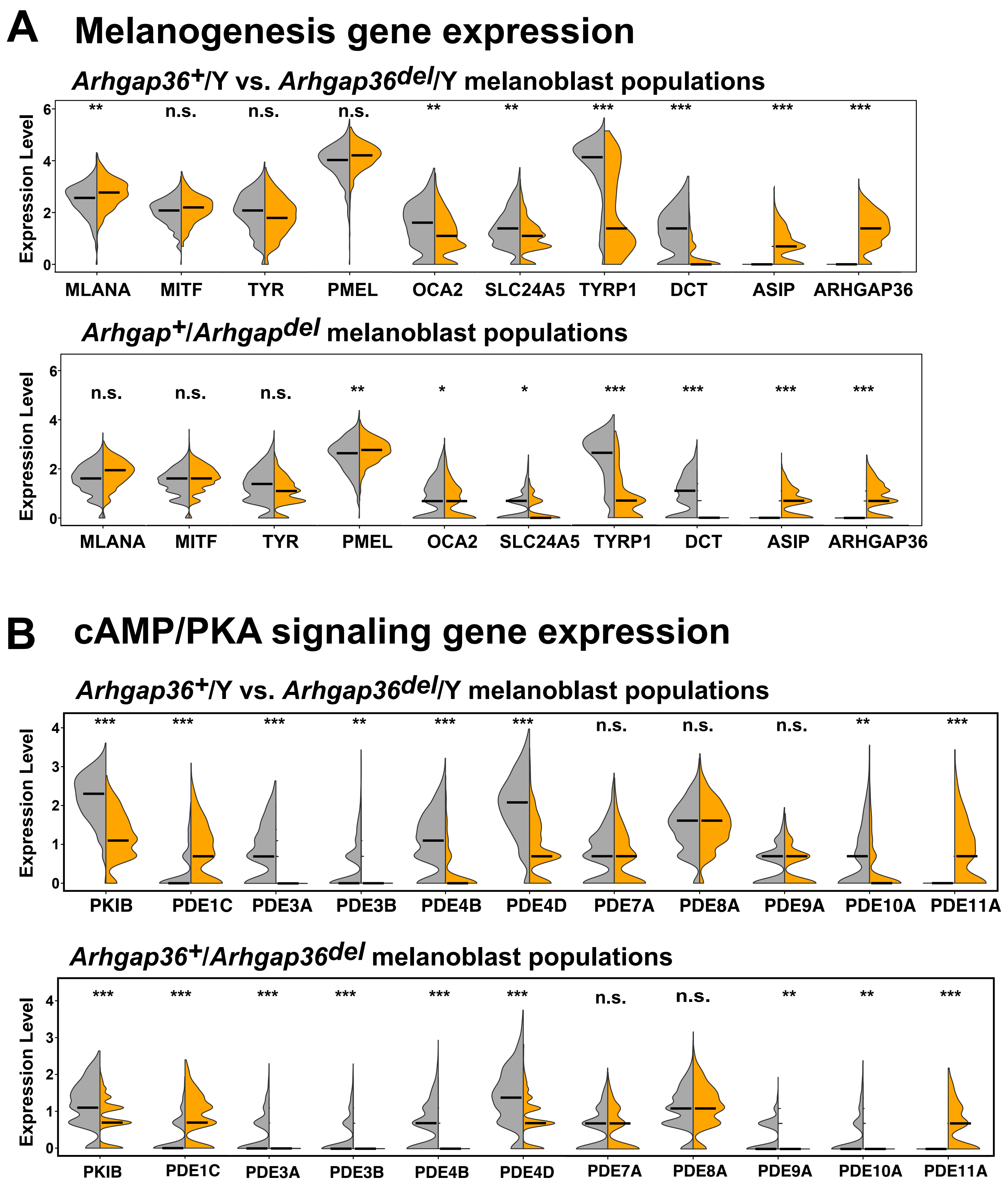

### Supplementary Figure 4

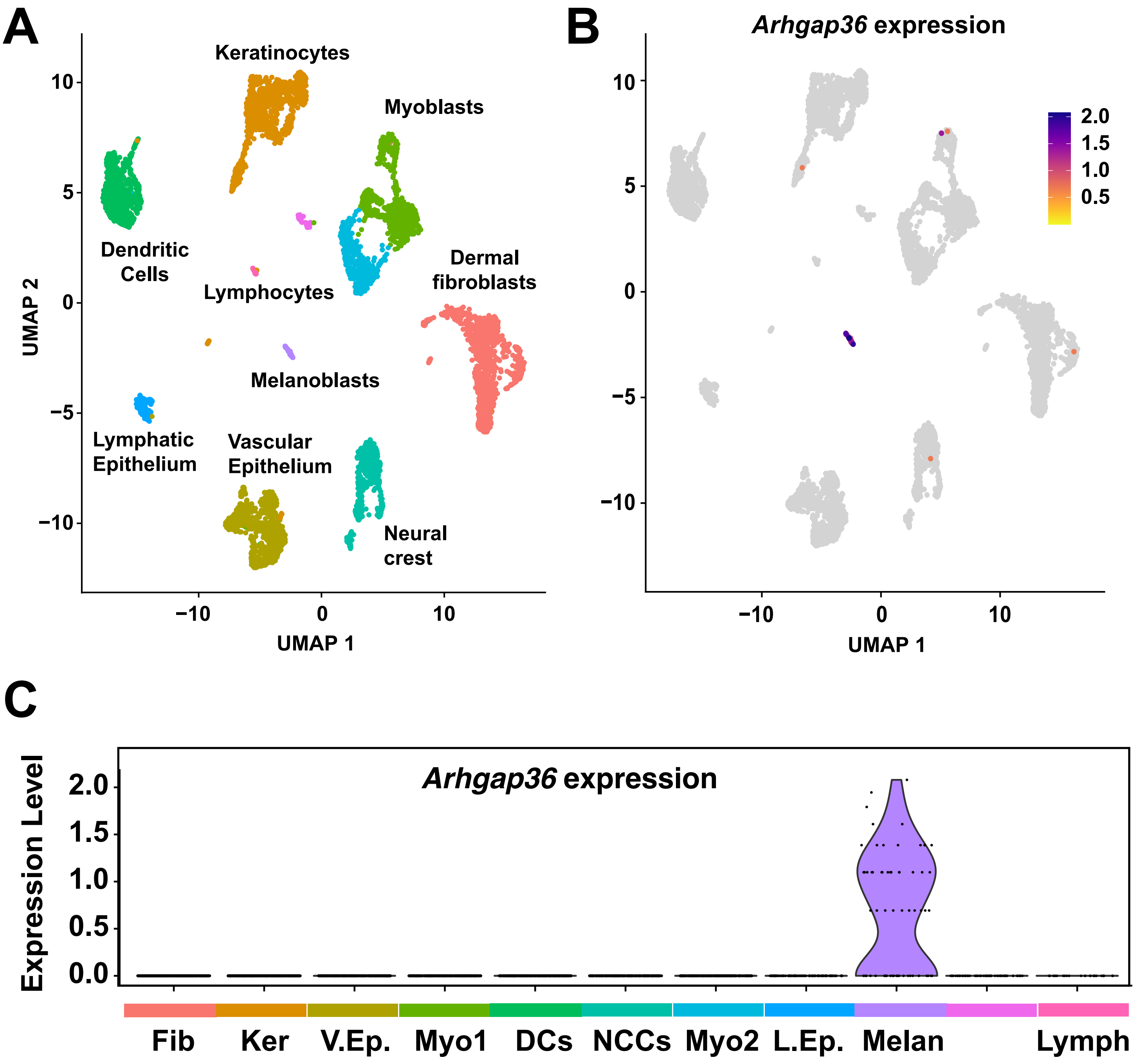

### Supplementary Figure 5

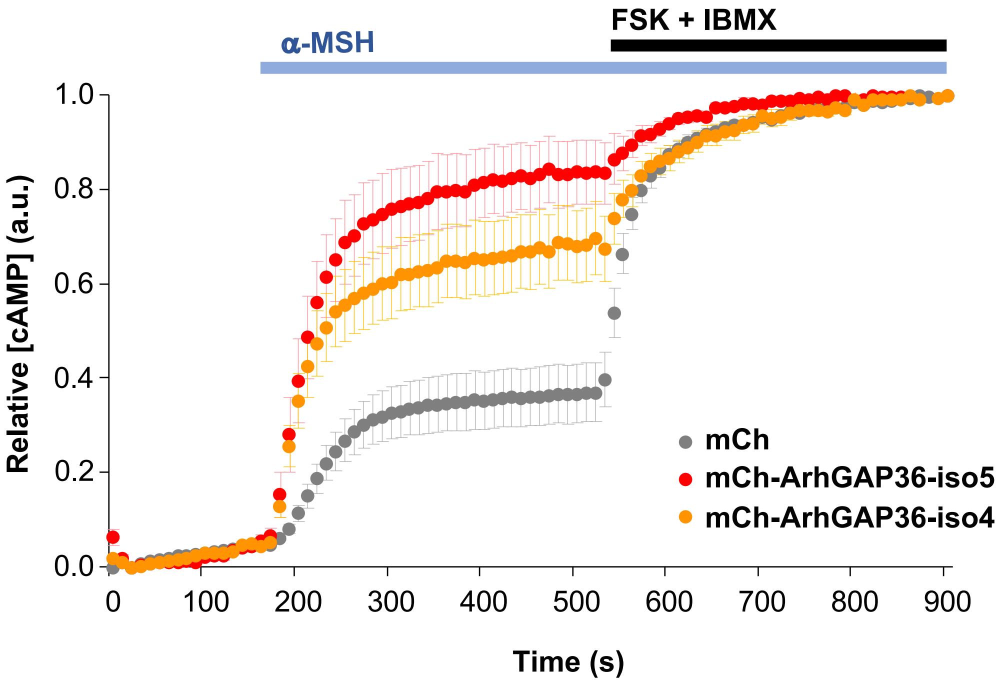

### Supplementary Figure 6

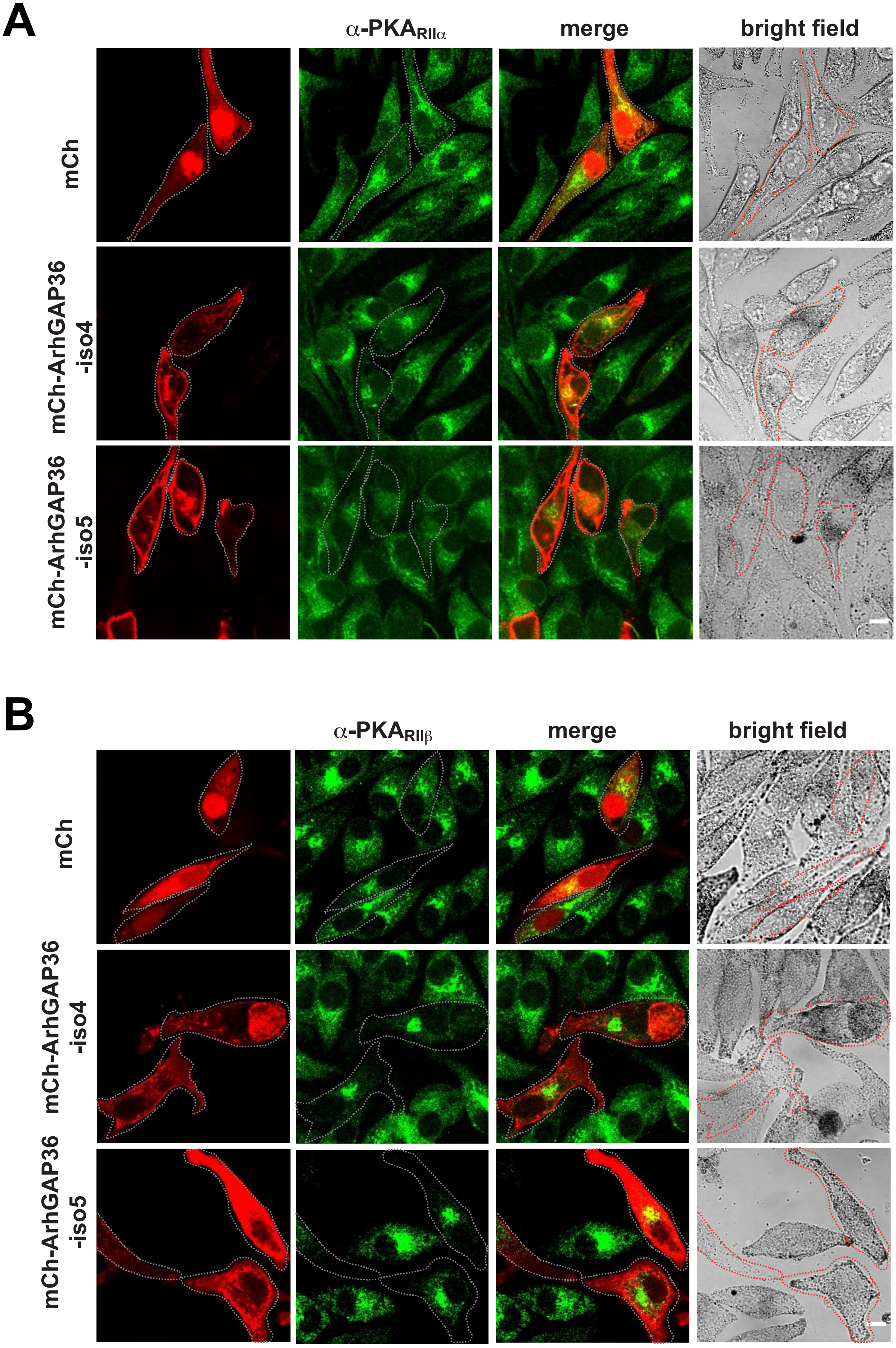
